# Virally-Mediated Enhancement of Efferent Inhibition Reduces Acoustic Trauma in Wild Type Murine Cochleas

**DOI:** 10.1101/2024.09.12.612688

**Authors:** Eleftheria Slika, Paul A. Fuchs, Megan Beers Wood

## Abstract

Noise-induced hearing loss (NIHL) poses an emerging global health problem with only ear protection or sound avoidance as preventive strategies. In addition, however, the cochlea receives some protection from medial olivocochlear (MOC) efferent neurons, providing a potential target for therapeutic enhancement. Cholinergic efferents release ACh (Acetylycholine) to hyperpolarize and shunt the outer hair cells (OHCs), reducing sound-evoked activation. The (α9)_2_(α10)_3_ nicotinic ACh receptor (nAChR) on the OHCs mediates this effect. Transgenic knock-in mice with a gain-of-function nAChR (α9L9’T) suffer less NIHL. α9 knockout mice are more vulnerable to NIHL but can be rescued by viral transduction of the α9L9’T subunit. In this study, an HA-tagged gain-of-function α9 isoform was expressed in wildtype mice in an attempt to reduce NIHL. Synaptic integration of the virally-expressed nAChR subunit was confirmed by HA-immunopuncta in the postsynaptic membrane of OHCs. After noise exposure, α9L9’T-HA injected mice had less hearing loss (auditory brainstem response (ABR) thresholds and threshold shifts) than did control mice. ABRs of α9L9’T-HA injected mice also had larger wave1 amplitudes and better recovery of wave one amplitudes post noise exposure. Thus, virally-expressed α9L9’T combines effectively with native α9 and α10 subunits to mitigate NIHL in wildtype cochleas.

**One Sentence Summary:** Viral transduction of a gain-of-function nAChR enhances the native cholinergic inhibition to protect the cochlea from noise-induced hearing loss.

## INTRODUCTION

With the ever-accelerating use of personalized sound devices and urban exposure to high sound pressure levels, NIHL is rising as a major health problem worldwide. In their 2021 report, the World Health Assembly predicts that by 2050 almost 2.5 billion people will suffer from hearing loss[1, 2]. In 2019, 1.57 billion people suffered to some degree of hearing loss, equivalent to 43.45 million years lived with disability, making hearing loss the third largest cause of global disease burden. This corresponds to more than $981 billion in total costs, with most of this burdening populations from lower socioeconomic groups and older age[3-5]. Noise exposure, either occupational or social, accumulated over one’s lifespan, promotes sensory hair cell damage and loss of auditory function[6]. Aside from protective devices and avoidance of loud sounds, there are no FDA approved therapies to prevent noise-induced damage to the ear. Here, we will describe how inner ear gene therapy can enhance a native reflex of the cochlear apparatus to mitigate noise-induced hearing loss in vivo. This approach is a significant advance for gene therapy; not to replace a defective gene, but to enhance the function of a native neuronal circuit by addition of a gain-of-function isoform.

The medial olivocochlear reflex provides efferent innervation from the superior medial olivary complex in the brainstem to the sensory epithelium of the cochlea, the organ of Corti [7, 8]. Prior to hearing onset in mammals, cholinergic medial olivocochlear fibers synapse with IHCs, contributing to the differentiation and functional maturation of the IHC afferent synapses [9-11]. As the inner and outer hair cells differentiate, medial olivocochlear fibers migrate from the inner to the outer hair cell region, where they form inhibitory synapses on OHCs. By hearing onset, the medial efferent fibers synapse only with the OHCs. The medial olivocochlear locus in the brainstem receives input from bilateral cochlear nuclei to activate efferent fibers in response to elevated sound levels. The pharmacology of the medial olivocochlear synapse involves the (α9)_2_(α10)_3_ nicotinic receptor, which is activated by the binding of 2 acetylcholine molecules on the α9 subunits [12-14]. Upon medial efferent activation, ACh release binds the nAChR and induces an inward Ca^2+^ current, which, in turn, activates SK and BK Ca^2+^-dependent K^+^ channels [15, 16]. This increase in membrane permeability to K^+^ hyperpolarizes the OHC and shunts their electromotility, thus preventing noise-induced overactivation.

The medial olivocochlear system acts with frequency and sound level specificity, to fine tune OHC electromotility and prevent non-specific excitation. Several studies have shown that it can act as a protective mechanism from noise-induced hair cell damage [17-28]. Substituting a threonine for a leucine at position 9’ (L9’T) of the second transmembrane domain of the α9 subunit produces an α9L9’T isoform with increased sensitivity to ACh and prolonged mean open times of the channel. This results in enhanced magnitude and duration of the MOC effect [29]. Studies on transgenic knock-in mice for the gain-of-function α9L9’T isoform showed that they were better protected from noise-induced threshold shifts than the wild type (control) group for the same noise exposure. In contrast, transgenic α9 knockouts, lacking MOC feedback, were more vulnerable to noise-induced threshold shifts [17]. Viral transduction of the α9L9’T isoform in α9 knockout mice spared them from the temporary threshold shifts that their uninjected α9 knockout littermates suffered after noise exposure [28]. Fluorophore-conjugated conopeptides showed expression of the α9L9’T nAChR on the synaptic membrane of α9 knockout OHCs.

AAV2.7m8 efficiently transfects all types of neuroepithelial cells in the cochlea, including both inner and outer hair cells [30]. For visualization of the α9 subunit on fixed tissue through immunolabeling, an HA-tag was attached to the C-terminal of the α9 sequence. This HA-tag position was shown previously not to reduce receptor function in α9-HA heterozygous mice [31]. In this study, the α9L9’T-HA sequence (expressed under a CAG promoter in AAV2.7m8-Figure 1A) is introduced into wild type murine (C57BL/J6) cochleas through posterior semicircular canal injections. The aim is to express the α9L9’T-HA subunit on the OHC postsynaptic membrane for interaction with native α9 and α10 to form functional pentameric nAChRs, thus enhancing MOC feedback to increase protection against noise-induced trauma.

**Fig. 1.**
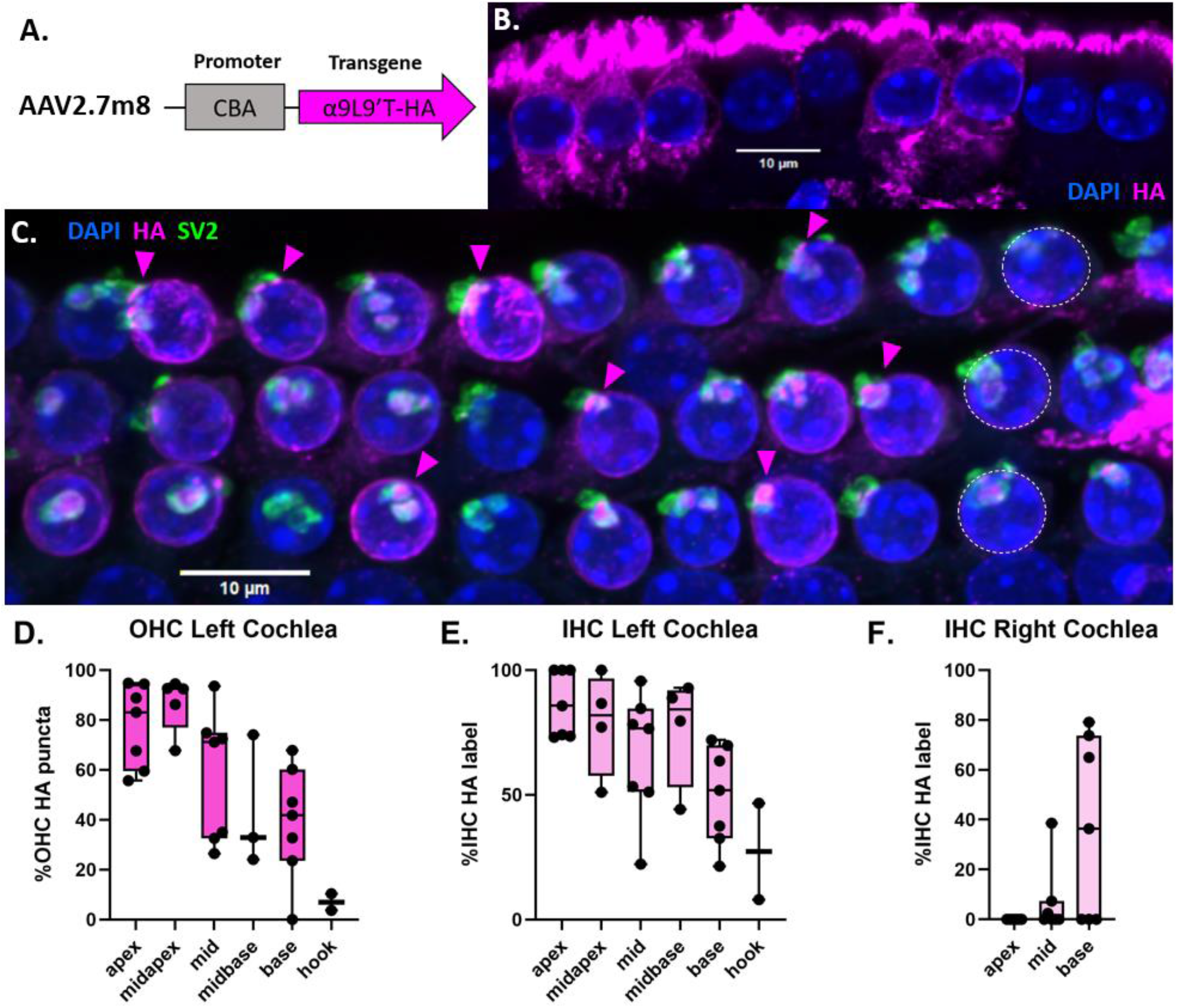
Viral transduction of cochlear α9L9’T-HA expression. **A**. Schematic of AAV2.7m8 bearing the α9L9’T-HA sequence under a strong universal promoter.**B** IHC row of the left basal turn of a 7-week-old C57Bl/J6 wild type male mouse injected at P3 with 500 nl of AAV2.7m8-CAG-α9L9’T-HA viral solution. IHCs with HA cytoplasmic label in magenta and nuclei stained in blue with DAPI. **C**. Maximal intensity projection image of the 3 OHC rows (1 nucleus in each row indicated by white dashed lines) of the left middle turn of a 7.5-week-old C57Bl/J6 wild type female mouse injected at P3 with 900nl of AAV2.7m8-CAG-α9L9’T-HA viral solution. Immunofluorescent secondary antibodies label dense HA puncta (magenta, some indicated by arrow heads) on the OHC postsynaptic membrane juxtaposed to SV2-positive (green) presynaptic terminals, showing synaptic localization of the virally encoded α9 subunit. OHC and supporting cell nuclei are labeled with DAPI (blue) **D-F**. HA expression gradient along the cochlear spiral. **D**. Percent (mean ± max, min and individual values) of OHC with postsynaptic HA expression in 7 left (injected) cochleas; apex = 7 segments, 847 OHCs, mid apex 5 segments, 717 OHCs, mid 7 segments, 903 OHCs, mid base 3 segments, 459 OHCs, base 7 segments, 794 OHCs, hook 2 segments, 197 OHCs. **E**. Left (injected) cochlea (N=7) percent (mean ± max, min and individual values) of IHC with HA cytoplasmic label, apex = 7 segments, 199 IHCs, midapex 4 segments, 157 IHCs, mid 7 segments, 257 IHCs, mid base 4 segments, 179 IHCs, base 7 segments, 271 IHCs, hook 2 segments, 70 IHCs. **F**. Percent of IHC (mean ± max, min and individual values) with HA cytoplasmic label in 7 right cochleas, 675 IHCs.

## RESULTS

### 1. α9L9’T-HA is expressed on the postsynaptic membrane of OHCs

Immunofluorescent labeling of the HA tag of the virally introduced gain-of-function α9 subunit showed widespread expression along the cochlear spiral (magenta - Figure 1C). This was combined with presynaptic labeling of the efferent fibers by an SV2 antibody (green, Figure 1C), along with DAPI which labels cell nuclei (blue, Figure 1C). HA puncta juxtaposed to SV2-positive efferent terminals were found on the basal pole of OHCs, near the nucleus. This corresponds to the known efferent contacts on OHCs[7, 8, 12-15, 32] indicating that the viral protein is localized as expected for the α9α10 nAChR. Quantification of the fraction of OHCs with HA synaptic puncta in confocal images from each cochlear region (Figure 1D) showed a declining gradient from apex to base in the injected (left) cochleas. Starting at the apical turn, the percentage of HA-positive OHCs declined from 78% to just 7% in the basal-most hook.

Synaptic HA puncta (and cholinergic responses) are absent from adult IHCs[9] but there was substantial diffuse cytoplasmic HA immunolabel. The high AAV2.7m8 transfection efficiency and strong CAG promoter were shown previously to drive expression of viral protein in the cytoplasm of IHCs and supporting cells[30], confirming successful injection, viral transfection and patterns of expression for these preparations. As with OHC expression, the IHC cytoplasmic-HA label declined from apex to base, from 87% in the apex to 27% in the basal-most hook region (Figure 1E).

Limited HA label also was observed in the contralateral (uninjected) cochlea. In contrast to the injected side, HA labeling was confined mainly to the basal regions of the uninjected ear. Cytoplasmic HA label declined steeply from 36% of IHCs at the base, to just 7% at the middle turn and none at the apex (Figure 1F). Virtually no OHC postsynaptic HA puncta were seen in the contralateral cochleas, except for one OHC in the basal segment of each of two cochleas.

### 2. Baseline ABR thresholds and wave1 amplitudes of the α9L9’T-HA vs the uninjected control group are slightly elevated

Viral injections were made into the left posterior semicircular canal of C57Bl/J6 mouse pups at postnatal days 2-4 (P2-4). The experimental group included 19 α9L9’T-HA injected C57Bl/J6 wild-type mice. The control groups included 14 uninjected and 8 GFP-injected littermates.

Baseline ABRs (before noise-exposure) were recorded from the left (injected) side of 5-week-old animals for clicks and pure tones at 8, 12, 16, 24, 32, 40 and 46 kHz. Pairwise comparison of click and pure tone thresholds found no significant differences between α9L9’T-HA injected and control groups (Kruskal Wallis test for clicks, non-normal distribution; multiplicity adjusted p-values with Tukey’s multiple comparisons test after two-way ANOVA for pure tone frequencies - Figure 2A). However, two-way ANOVA did show significance between groups (F (2, 266)= 5.165, **p<0.01) for pure tones. The mean difference in threshold between α9L9’T-HA and uninjected groups was 6 dB (95% CI 2-10 dB; **p<0.01). But, mean thresholds did not differ significantly between the α9L9’T-HA and the GFP-injected groups (1dB, 95% CI -4 - 7dB) or between the uninjected and GFP-injected groups (−4dB, 95% CI -10 – 1dB). (Multiplicity adjusted p-values with Tukey’s multiple comparisons test).

**Fig. 2.**
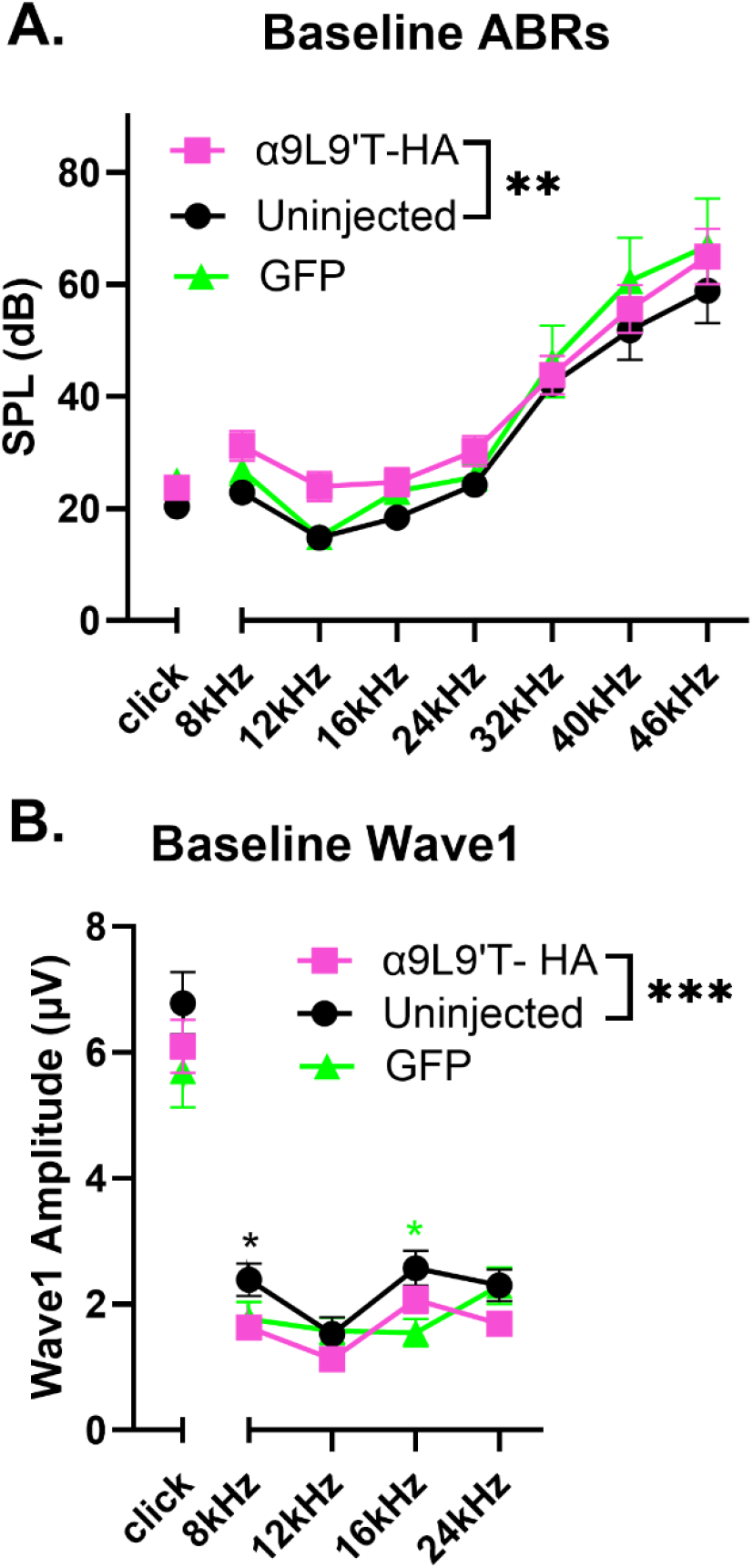
Baseline ABR thresholds and click wave1 amplitude. ABR recordings pre-exposure for all the groups (mean ± SEM). C57Bl/J6 wild type mice 5 weeks old. α9L9’T-HA injected group N= 19 (magenta) vs the uninjected group N=14 (black) vs GFP-injected group N= 8 (green). Black asterisks for uninjected vs α9L9’T-HA, magenta asterisks for GFP vs α9L9’T-HA, green asterisks for GFP vs uninjected. **A**. ABR thresholds at baseline. Kruskal Wallis test for click thresholds and two-way ANOVA for pure tones with multiplicity adjusted P-values after Tukey’s multiple comparisons test. **B**. 80dB wave1 amplitudes at baseline. Ordinary one-way ANOVA for clicks and two-way ANOVA for pure tones with multiplicity adjusted p-values after Tukey’s multiple comparisons test. *p<0.05, **p< 0.01, ***p<0.001, ****p<0.0001, ns non-significant.

Baseline click wave1 amplitudes at 80dB were not significantly different among the groups (ordinary one-way ANOVA – Figure 2B). However, 2-way ANOVA for amplitudes of pure tones up to 24 kHz did show significance among groups (F (2, 152) = 7.041, **p<0.01). As for thresholds, only α9L9’T-HA versus uninjected overall mean difference (−0.57 μV) is significant, whereas the α9L9’T-HA vs GFP difference (−0.169μV) and the uninjected vs GFP (−0.4μV) are not (multiplicity adjusted p-values with Tukey’s multiple comparisons test). Pairwise comparisons for pure tone wave1 amplitudes showed significance between the α9L9’T-HA and the uninjected group at 8kHz and between the uninjected and GFP-injected at 16kHz.

These baseline data suggest a slight threshold elevation due to expression of α9L9’T-HA, as reported previously ([29]), but this was evident only from 8 to 16 kHz. Thresholds for the highest frequencies tested were slightly, but not significantly, elevated for both α9L9’T-HA and GFP-injected mice. Likewise, baseline wave 1 amplitudes were minimally affected by viral injection. In contrast, viral injection markedly altered the response to acoustic trauma.

### 3. Post-acoustic trauma, the α9L9’T-HA group has lower thresholds & threshold shifts compared to controls

All animals were exposed at 5 weeks of age to an 8-16 kHz octave band signal at 100 dB for 1 hour (gray band in Figure 3). Post-exposure ABR threshold values and shifts from baseline were plotted for clicks and individual frequencies (8, 12, 16, 24, 32, 40 & 46 kHz) at 1-, 7-, and 14-days post-trauma (Figure 3). For all time points, two-way ANOVA showed significantly lower absolute thresholds and threshold shifts for the α9L9’T-HA group compared to GFP-injected and uninjected controls (Figure 3). GFP-injected and uninjected ears showed similar patterns of acoustic damage. In contrast to the two control groups, the α9L9’T-HA injected group not only had one day post trauma thresholds significantly lower than the controls, but there was also better recovery at 7 and 14 days after trauma. This is shown by the threshold shift graphs (Figure 3), which present a pattern similar to that of absolute thresholds. α9L9’T-HA injected shifts were the smallest and GFP-injected shifts were not different from those of the uninjected mice.

**Fig. 3.**
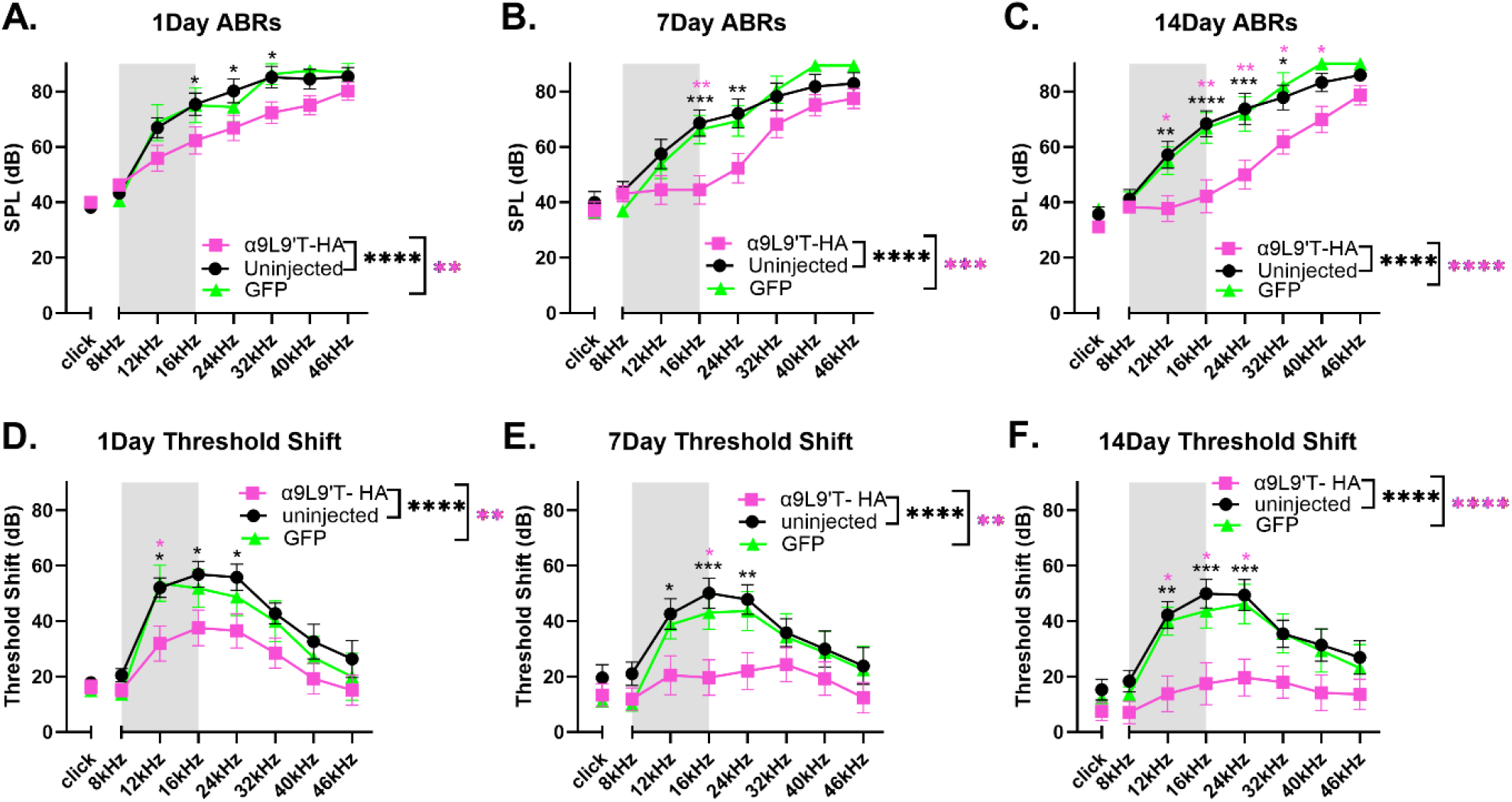
Post-exposure ABR absolute thresholds and threshold shifts from baseline (mean ± SEM) All animals were exposed to an octave-band (8-16 kHz) stimulus at 100 dB for 1 hour, represented by the gray area on the frequency band. α9L9’T-HA injected group N=19 (magenta), uninjectd group N=14 (black), GFP-injected group N= 8 (green). Black asterisks for uninjected vs α9L9’T-HA, magenta asterisks for GFP vs α9L9’T-HA, green asterisks for GFP vs uninjected. *p<0.05, **p<0.01, ***p<0.001, ****p<0.0001, ns non-significant. Ordinary one way ANOVA (if normality present) or Kruskal Wallis test (if non normal data) for clicks, no significance at any time point. Two-way ANOVA for pure tones with multiplicity adjusted p-values with Tukey’s multiple comparisons test for overall group means and individual frequency comparisons.

Additional features were revealed by within group comparisons (Figure 4). The α9L9’T-HA injected group recovered progressively from 1 to 14 days post trauma. Indeed, at 14 days Post-exposure the click, 8, 12, 16, 40 & 46 kHz absolute thresholds did not differ significantly from the pre-exposure baseline (Figure 4A). Thus, the α9L9’T-HA injected group experienced a smaller (mean 15dB above baseline) threshold shift (TS) compared to controls. The 12-24kHz area in particular showed the maximal difference between the α9L9’T-HA and the control groups initially, with better recovery as time progressed. Threshold shifts at 1, 7 and 14 days were progressively smaller and significantly different from each other for pure tones and clicks (Figure 4D). Repeated measures two-way ANOVA for pure tone threshold shifts (graphs 4D-F) with Tukey’s multiple comparisons for overall group means yielded significant p-values between 1 and 7 day for the uninjected and the GFP -injected animals, indicating partial recovery at 7 days. However, there was no further improvement at 14 days, suggesting a degree of permanent threshold shift (PTS). At 14 days, the mean threshold elevation was 36dB for the uninjected and 33dB for the GFP-injected group. Click threshold shift comparisons among 1, 7 and 14 day values were not significant for both groups (Friedman’s test and Dunnett’s test for multiple comparisons), emphasizing that there was minimal threshold recovery after the initial 1 day damage.

**Fig. 4.**
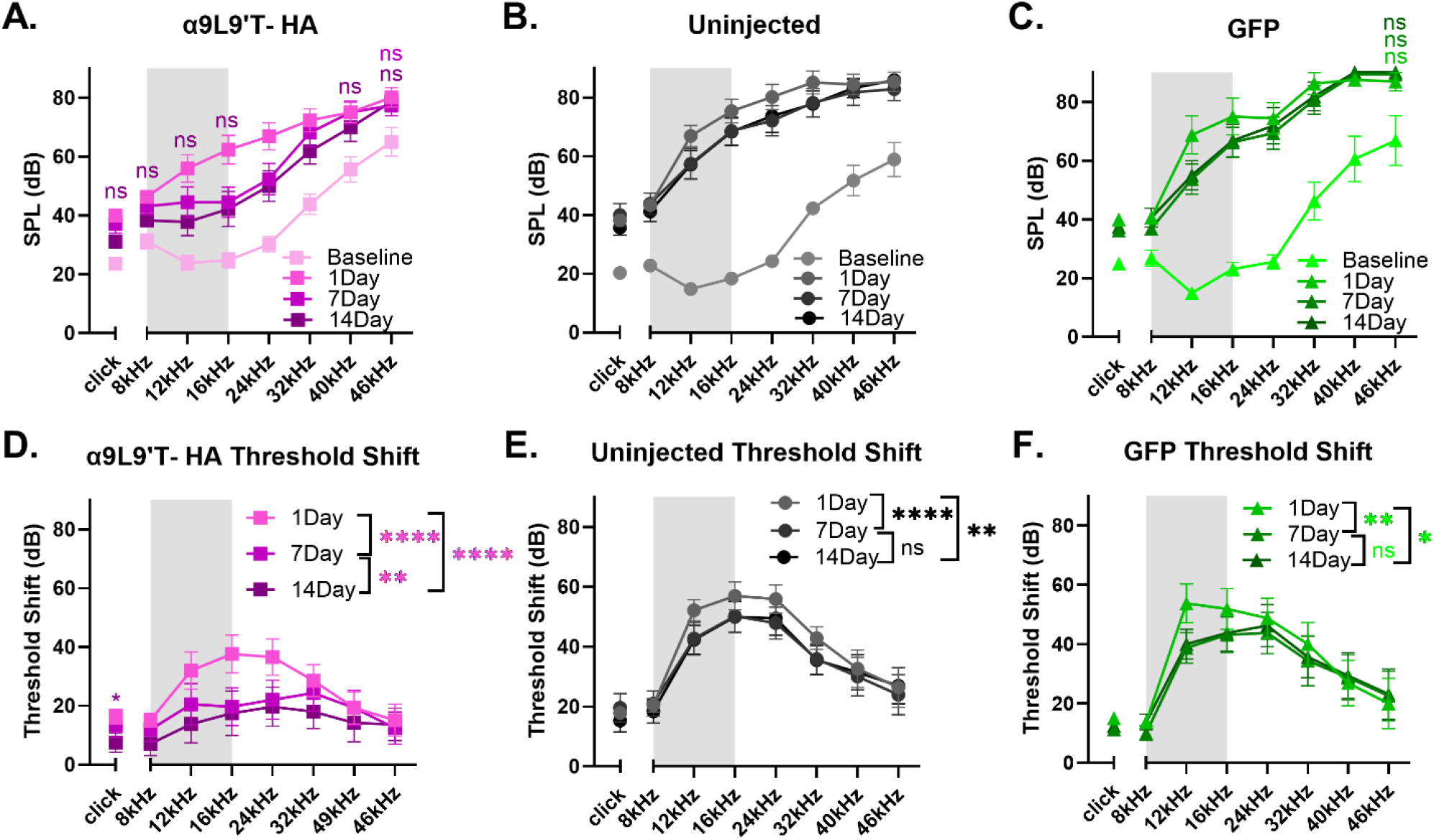
Threshold absolute values (A) and shifts from baseline (B) at 1, 7 and 14 days Post-exposure per group. (mean ± SEM) All animals were exposed to an octave-band (8-16 kHz) stimulus at 100 dB for 1 hour, represented by the gray area on the frequency band. α9L9’T-HA injected group N=19 (magenta), uninjected group N=14 (black), GFP-injected group N= 8 (green). Different color intensities represent different time points and same color asterisks compare Post-exposure time points to baseline of that group. non-significant (ns) points are depicted as indicators of protection or recovery from acoustic damage. **A-C**. For clicks, Friedman’s test with Dunn’s multiple comparisons test, for pure tones, repeated measures two-way ANOVA with Dunnett’ s multiple comparisons test. **D-F**. For clicks, Friedman’s test with Dunn’s multiple comparisons test, for pure tones, repeated measures two-way ANOVA with Tukey’ s multiple comparisons test. *p<0.05, **p<0.01, ***p<0.001, ****p<0.000, ns non-significant.

### 4. Post trauma, click and pure tone wave 1 amplitudes at 80 dB are largest for the α9L9’T-HA group

Wave 1 amplitude for clicks and pure tones was measured for all mice prior to and then 1, 7- and 14-days after acoustic trauma (Figure 5). α9L9’T-HA click amplitudes at 80 dB were significantly larger than controls at 14 days (Figure 5A-C & Figure 6), indicating that the α9L9’T-HA injections resulted in click amplitude recovery, which the control groups lacked (ordinary one way ANOVA, with Tukey’s multiple comparisons test). For all groups, click wave1 amplitude was halved 1-day Post-exposure. For the control groups, click amplitudes did not recover and remained significantly lower, around half of baseline values (Figure 5D-F, repeated measures one-way ANOVA with Dunnett’s multiple comparisons between group means, multiplicity adjusted p-values). For the uninjected mice, baseline mean at 14 days was 3.21μV, vs 6.78μV pre-trauma. For the GFP group, amplitude fell from 5.7μV to 2.93μV at 14 days. However, the α9L9’T-HA group showed excellent recovery, with 14-day click amplitude mean of 5.13μV not significantly different from baseline (6.1μV) (Figure 5D).

**Fig. 5.**
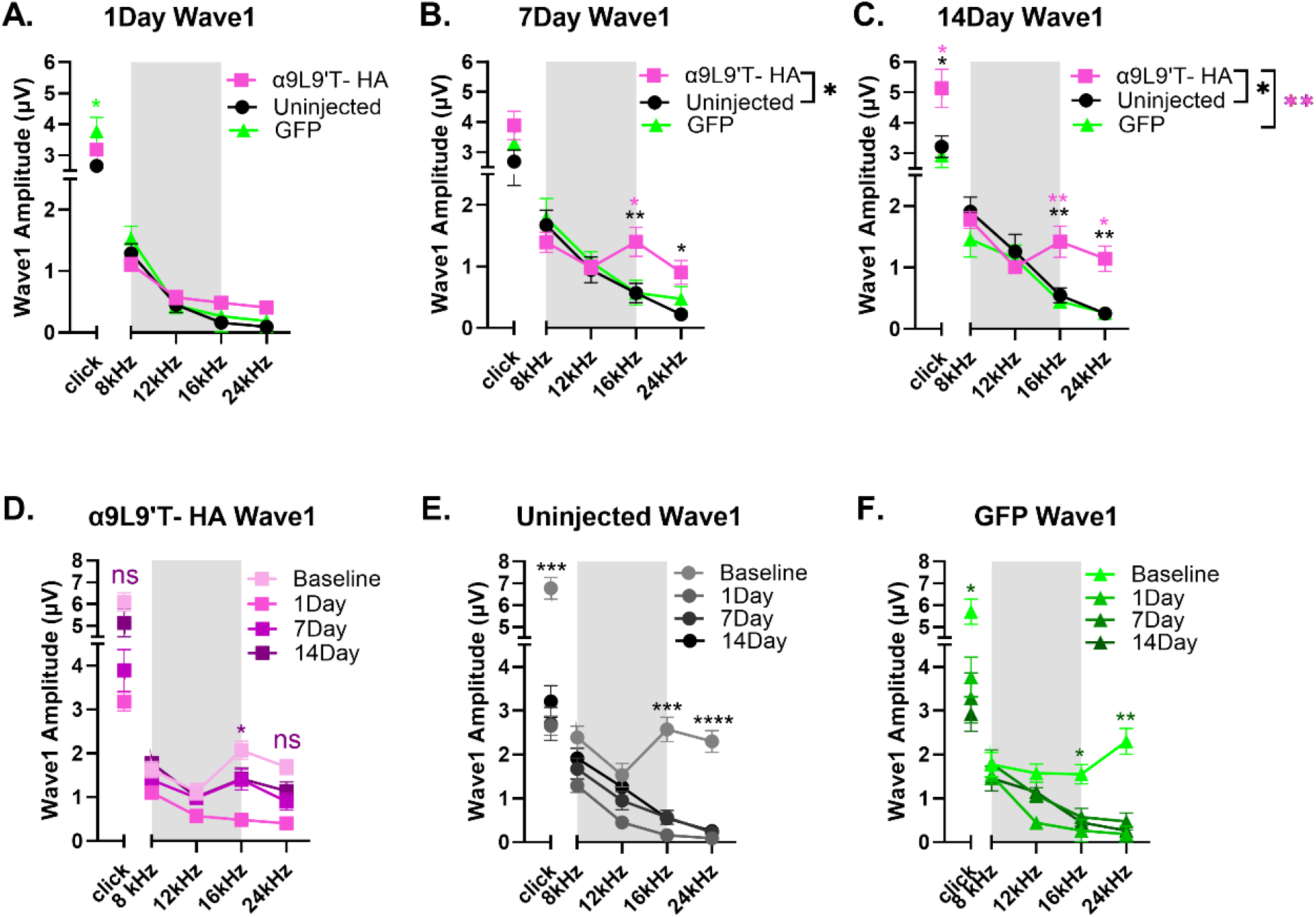
ABR wave 1 amplitude (μV) for clicks and pure tones. mean± SEM. All animals were exposed to an octave-band (8-16 kHz) stimulus at 100 dB for 1 hour, represented by the gray area. α9L9’T-HA injected group N=19 (magenta), uninjected group N=14 (black), GFP-injected group N= 8 (green). **A-C**. 1, 7 and 14 days Post-exposure comparisons between groups. Black asterisks for uninjected vs α9L9’T-HA, magenta asterisks for GFP vs α9L9’T-HA, green asterisks for GFP vs uninjected. **D-F**. Within group comparisons. Different color intensities represent different time points and same color asterisks compare Post-exposure time points to baseline of that group. Non-significant (ns) points are indicators of protection or recovery from acoustic damage. Multiplicity adjusted p-values of 14day vs baseline comparisons for clicks, 16 and 24kHz are plotted. *p<0.05, **p<0.01, ***p<0.001, ****p<0.0001, ns non-significant.

**Fig. 6.**
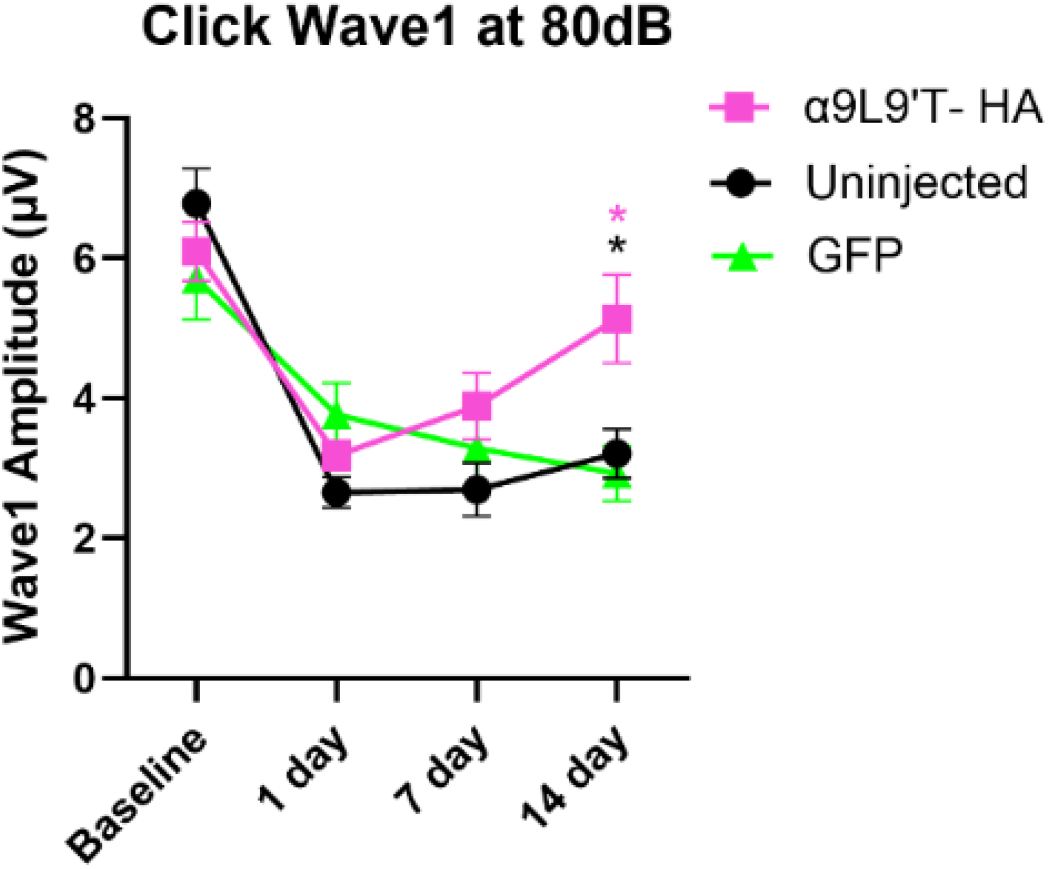
Click wave1 amplitude recovery over time. mean± SEM. Repeated measures two-way ANOVA with Tukey’s multiple comparisons test showed that at 14 days, the α9L9’T-HA group was significantly higher than the controls, showing progressive recovery, with no significant difference between pre-exposure and 14 day values.

In a similar pattern, pure tone wave 1 amplitude analysis of group means showed that α9L9’T-HA amplitudes were significantly higher than the uninjected at 7 & 14 days. Especially at 14 days, amplitudes for this group were higher than both control groups (two-way ANOVA with Tukey’s multiple comparisons test). With individual frequency comparisons (multiplicity adjusted p-values with Tukey’s test), the 16 & 24kHz frequencies showed larger amplitudes and maximal recovery for the α9L9’T-HA group than for controls (Figure 5A-C). Within group analysis showed that one day after acoustic trauma, wave 1 amplitude was reduced significantly in all cohorts. On subsequent days the α9L9’T-HA injected mice showed improvement. Wave 1 amplitude at 7 days and 14 days Post-exposure increased gradually from the low 1-day values for frequencies above 12 kHz, returning almost back to baseline at 14 days. In contrast, wave 1 amplitude of both GFP-injected and uninjected mice had no significant recovery 14 days post trauma (Figure 5D-F & Figure 6).

Of note, the 8 to 12 kHz frequency band was not affected by the octave-band exposure, thus wave 1 amplitudes of these tones remained the same at all time points for all groups. Similarly to ABR thresholds, maximal damage was located at the 16-24 kHz area, with only the α9L9’T-HA group showing amplitude recovery at 14 days.

### 5. IHC synapse counts show no significant difference in the α9L9’T-HA injected cochleas vs controls Post-exposure

After concluding ABR recordings, the animals were sacrificed at 15-17 days post-noise exposure and the synaptic puncta of IHC type I afferents were analyzed. For visualization of IHC afferent synapses, immunofluorescent labelling of presynaptic ribbons (anti-CtBP2) and postsynaptic densities (anti-PSD95) was performed. Confocal microscopy images from the apical, middle, and basal turns of each cochlea were acquired, with 15-30 IHC nuclei counted in each image. The total number of juxtaposed CtBP2/ PSD95 puncta in each image was counted and divided by the number of IHC nuclei. Synapses/IHC are plotted for the apical, middle, and basal cochlear region of the α9L9’T-HA injected left cochleas (N=3, magenta) and their contralateral right cochleas (N=3, pink) and compared with the left cochleas of uninjected (N=6, grey) and GFP-injected mice (N=8, green) (Figure 7). Two-way ANOVA showed no significant interaction between the 4 groups (F (6, 48) = 1.45, P>0.05). This observation could be attributed to the small number of mice studied. If validated in a larger group of animals, it could mean that this acoustic protocol does not necessarily cause quantifiable IHC synapse loss in these animals at this time point, but instead permanent IHC synapse dysfunction. Other structures in the organ of Corti could also be damaged, affecting the activation of the afferent cochlear fibers. Whatever the cause of these observations, IHC synaptic puncta counts failed to detect a difference among these groups. Nevertheless, even if synapse counts have recovered after trauma, they might not function properly and remain dysregulated.

**Fig. 7.**
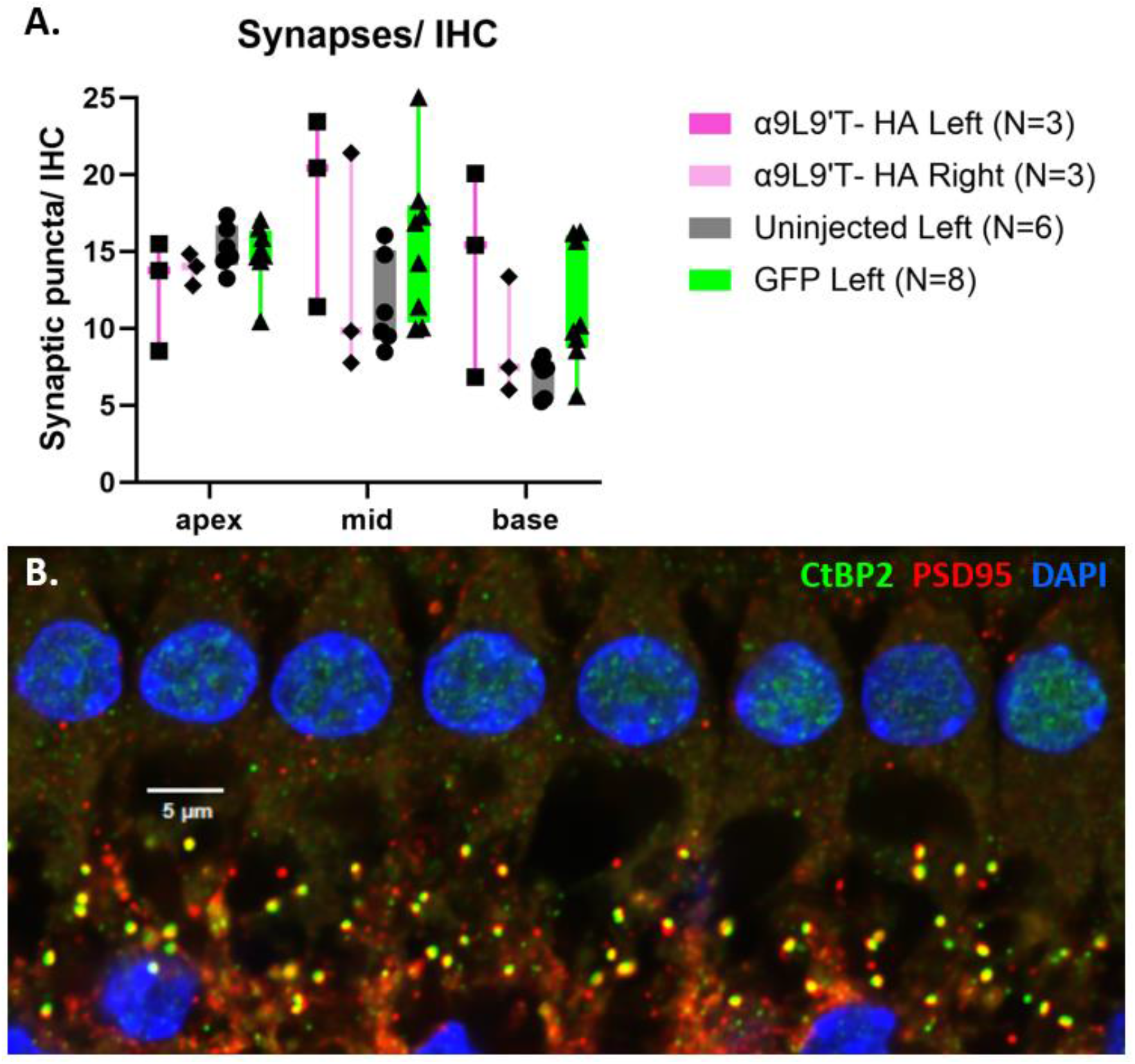
IHC synapse counts. **A** Synapses/ IHC for the apical, middle and basal area of the left α9L9’T-HA injected cochleas, the right uninjected α9L9’T-HA side and the left cochleas of the uninjected and GFP-injected groups. Confocal images of every cochlear region, with 15-30 IHC per image, were acquired. Co-localized immunolabelled CtBP2/ PSD95 puncta counted and divided by the number of IHC nuclei stained with DAPI. Mean ± max, min and individual values. Two-way ANOVA showed no difference between groups in all cochlear regions **B** Representative image of co-localized CtBP2 (green)/ PSD95 (red) synaptic punctae from the left middle turn of a GFP-injected 7 week old male C57Bl/J6 mouse. IHC nuclei are stained blue with DAPI.

## DISCUSSION

Previous work had shown that heterozygous α9L9’T knock-in mice had greatly enhanced efferent effects such as suppression of DPOAEs and acoustic protection from noise exposure compared to wildtype[29], raising the possibility of combined, functional wildtype and α9L9’T nAChRs. On that basis, the present studies were undertaken to determine if virally expressed nAChR subunits could insert significantly into wildtype synaptic complexes. And indeed, such integrated expression does occur, as shown both by immunolabeling of an HA-tag, and by functional measures (acoustic protection).

α9L9’T-HA is localized to the OHC postsynaptic membrane after AAV2.7m8 transfection with the transgene under a strong CAG promoter[30, 33]. On the injected side (left), postsynaptic HA puncta in OHCs, and cytoplasmic HA immunolabel of IHCs was greatest in apical segments and declined toward the base. This pattern would result if virus could only access hair cells in the organ of Corti via the scala tympani, but not from endolymph across tight junctions of the reticular lamina. As shown by dye tracking studies [32], injection into the posterior semicircular canal spreads through perilymph to scala vestibuli and the cochlear helicotrema, entering the scala tympani at the cochlear apex. Only then can virus access the hair cell membrane via fluid and electrolyte exchange between scala tympani and the organ of Corti. Thus, apical hair cells see the highest concentration of virus. Diffusional arguments also explain the opposite HA-expression gradient observed for the contralateral cochlea. Here, viral crossover from the injected side to the contralateral cochlea occurs through the aqueduct that drains the perilymph into the subarachnoid space and inserts close to the base of the cochlea, especially in mouse pups that have a patent aqueduct at this age [34-36]. Thus, the basal contralateral cochlea sees the highest concentration of virus [32].

At 1-, 7- and 14-days Post-exposure, both ABR absolute values and threshold shifts from baseline were significantly lower for the experimental α9L9’T-HA injected mice versus the control groups (Figure 3). The protective effect was greatest in the 12-32 kHz frequency range where maximal damage occurs adjacent to the noise band used for acoustic trauma [37-39]. This corresponds to the middle cochlear turn where efferent innervation density is greatest and α9L9’T-HA subunit expression expected to have the greatest impact. With time after noise exposure, the α9L9’T-HA group showed accelerated recovery compared to control groups. This suggests that enhanced efferent feedback can slow the progression from temporary threshold shift (TTS) to permanent threshold shift (PTS). This is reminiscent of observations made in mouse lines where equivalent acoustic trauma produced only TTS in wildtype, but PTS in α9 knockout mice[17].

Click wave1 amplitudes in all groups declined one day after acoustic trauma, with minimal recovery after that for the control groups. At 14 days post trauma, the α9L9’T-HA group had prominent recovery of wave 1 amplitudes, reaching pre-trauma amplitude values, while the control groups remained reduced (∼50% of baseline values). For a saturating, loud, broadband, acoustic stimulus, wave 1 amplitude reflects the number of activatable type I cochlear afferents (viable IHC synapses)[40-46]. Presuming that a click at 80dB is saturating, α9L9’T-HA injection may have had a benefit for IHC synapse function as well. Pure tone amplitudes showed a similar pattern at 16-32 kHz, where the α9L9’T-HA group’s amplitudes recovered over time, while the controls’ amplitudes remained low.

To probe IHC function further, ribbon synapses were counted 15-17 days after acoustic trauma. However, no statistically significant differences in synapse number were found as a function of experimental group. This is perhaps not surprising since synapse counts were quite variable. Although the absolute synapse numbers may not be significantly different, synaptic function and perhaps afferent fiber type distribution (high vs low spontaneous rate) may be different among the groups post-trauma. Noise-induced damage also targets other structures upstream of IHC afferent synapses, the stereocilia being one of them [47]. Such functional differences could contribute to the wave1 amplitude differences between the α9L9’T-HA group and that of the two control groups.

This work is motivated by the possibility of ‘efferent gene therapy’ as a therapeutic strategy. Is this feasible? Gene therapy for inner ear disease has advanced significantly, largely to correct ‘deafness genes’ that underlie inherited hearing loss. Notably, clinical trials for replacement, or editing of mutant otoferlin (DFNB9) are underway in several countries [48, 49]. Additional strategies to prevent or ameliorate ototoxic damage (as from antibiotics or chemotherapeutics) seek small molecules to address mitochondrial function, generation of reactive oxygen species, or aspects of calcium excitotoxicity. Prevention or reduction of noise-induced hearing loss however, remains largely the domain of protective coverings, or avoidance of acoustic trauma. Small molecules such as positive allosteric modulators are being considered to increase hair cell nAChR gating, thereby enhancing acoustic protection[50, 51]. In a study with heterologous cell plasmid transfection, choline acetyltransferase activity, directly leading to ligand binding regulation, proves a prerequisite for α9α10 nAChR assembly and membrane stabilization, while transmembrane proteins associated with hearing loss like TMIE or TIMEM132e serve as auxiliary subunits for channel gating [52]. Any future therapeutic must cross the blood/labyrinth barrier and avoid deleterious side effects. This may be a smaller problem due to the unique pharmacology and restricted expression of α9α10 nAChRs that have not been shown to function elsewhere in the nervous system[53]. However, a significant body of evidence has shown α9 expression in lymphocytes and implicated their activity in inflammatory pain [54-56]. Such concerns will be greater if regular dosing is required for protection from daily exposure to noise.

A gene therapy strategy offers some advantages. Direct application to the inner ear fluid space would greatly minimize, if not eliminate, side effects. If viral expression is sufficiently long-lasting there may be no need for repeated dosing. The drawbacks are that viral injection is time-consuming, requires surgical expertise, and is intimidating compared to an oral or injectable medication. Nonetheless, the risk/benefit profile might be compared favorably to that for cochlear implants where the surgical risks are higher, but the benefit well-established. Should efferent gene therapy be effective, the benefit for those at risk of early onset presbycusis and unavoidable noise exposure may justify the additional challenges. A second potential application is for those suffering painful hyperacusis, or noxacusis. If as proposed, type II cochlear afferents signal acoustic pain[57-60], enhanced efferent inhibition of glutamate release from OHCs onto type II afferents may be particularly beneficial for this condition.

## MATERIALS AND METHODS

### Study Design

The objectives of the study were to determine whether the virus would successfully transduce wild type murine cochleas with the α9L9’T-HA subunit. This was proved with immunohistological labelling of the HA-tag on the OHC membrane juxtaposed to SV2 presynaptic labelling (Figure 1). Subsequently, the functional implications of the gene therapy were studied by performing hearing assessment with ABR recordings. At baseline, the α9L9’T-HA subunit did not significantly affect auditory function (Figure 2). The primary goal was to study how the expression of the α9L9’T-HA subunit would affect the ABR response after traumatic exposure, compared to controls (Figures 3-6). Finally, IHC afferent synapses were labelled with immunofluorescent antibodies and counted to determine whether the observed findings would correlate with IHC afferent synapse counts at 14 days Post-exposure (Figure 7). Wild type C57Bl/J6 male and female mouse pups received posterior semicircular canal injections at postnatal days 2-5 (P2-P5) with either AAV2.7m8-CAG-α9L9’T-HA or AAV2.7m8-CAG-GFP virus. Uninjected littermates served as controls. The GFP-injected mice served as surgical controls. At 5 weeks of age all mice were exposed to an octave-band signal (8-16 kH)z at 100 dB SPL for 1 hour. Baseline ABRs were recorded 1-3 days prior to noise-exposure and 1, 7 and 14 days post noise-exposure. At 14 days of age, all mice were sacrificed and their cochleas fixed for immunohistological analysis.

This was a randomized experimental study. Treatment included posterior semicircular canal viral injection with AAV2.7m8-α9L9′T-HA.

Mice were purposely bred for this study, and breeding was carried out independently. Mouse pups were randomly and independently of any particular factor assigned to an experimental group. No power analysis was performed. Sample size was determined based on previous experience. 41 mice in total, 20 females and 21 males were used. The α9L9’T-HA group included 19 mice, 10 female and 9 male, the uninjected group included 14 mice, 6 female and 8 male and the GFP group included 8 mice, 4 female and 4 male.

The experiment is reported according to ARRIVE guidelines.

### α9L9’T-HA AAV2.7m8

Mouse α9L9′T-HA cDNA [31] was incorporated into a plasmid (Genewiz, South Plainfield, NJ 07080) and submitted to the Penn Vector Core (Gene Therapy Program, University of Pennsylvania School of Medicine) for incorporation into AAV2.7m8 [61] obtained from Addgene, University of California, Berkeley, MTA no. 486064. The resulting viral vector, (AAV2.7m8.CAG.mChrnα9L9′THA.bGH) was provided for use at 1.73 × 10^13^ viral copies/mL. This was subdivided into 100 μL aliquots and stored at -80°C. Once thawed for use, each aliquot was stored at 4°C (not refrozen) until exhausted. The same method was used for the GFP virus as employed previously [28, 30].

### Mice

Care and housing of animals was in accordance with institutional guidelines as specified in Johns Hopkins IACUC protocol MO23M04. C57Bl/J6 wild type mice were used for this study. The breeders were purchased from The Jackson Laboratory and maintained in the Johns Hopkins University School of Medicine Research Animal Resource facility. Mice were placed on a 12-h light-dark cycle and housed in cages with water and autoclaved feed, in a low-noise satellite facility without automated racks. All experiments were carried out under protocols approved by the Institutional Animal Care and Use Committee protocol #MO23M04.

### Animal Surgery

Hypothermia was used to induce and maintain anesthesia. The posterior semicircular canal was exposed through a postauricular incision and tissue dissection. Only the left ear of each animal was injected. A Nanoliter Microinjection System (Nanoject III, Model #3-000-207, Drummond Scientific Company, Broomall, PA) was used. 700-1100nl of viral solution (at 10^13^ GC/mL) mixed with Fast Green dye (4:1) were injected into the posterior semicircular canal using a glass micropipette (3.5”, Drummond Scientific Company #3-000-203-G/X) pulled to produce a tip opening of 20 μm. The pipette was backfilled with mineral oil and the injection solution loaded on it prior to the surgery. The solution was injected in bouts of 100nl at a rate of 50 nl/sec. Once the injection was concluded, the pipette was removed and the tissue and skin was sutured back together. The pups were put on a heating pad for recovery. The wound site was disinfected with betadine and 70% ethanol and the entire procedure carried out under sterile conditions.

### Auditory Brainstem Response

ABR measurements were conducted in a sound-treated booth (Industrial Acoustics Company, Bronx, NY; 59 × 74 × 60 cm^3^) lined with acoustic foam (Pinta Acoustic, Minneapolis, MN). Animals were anesthetized with intraperitoneal ketamine 100 mg/kg and xylazine 20 mg/kg and their eyes were swabbed with petrolatum-based ophthalmic ointment to prevent corneal ulcers. During the test they were placed on an infrared heating pad (Kent Scientific, Physio Suite, Modular System) to maintain a core temperature of 37 °C. Subdermal needle electrodes (Disposable Horizon, 13 mm needle, Rochester Med, Coral Springs, FL) were placed on the vertex (active), ipsilateral mastoid (reference), and hind limb (ground) in a standard ABR recording montage. Only the left ear was tested (random and counterbalanced selection). ABR signals were acquired with a Medusa4Z preamplifier (12kHz sampling rate) and filtered from 300–3000Hz with an additional band-reject filter at 60Hz. Clicks or pure-tone stimuli (512 repetitions 10 to 90dB in 10dB steps, 21 stimuli/s) were used to generate averaged ABR waveforms. The duration of the tonal stimulus was 5ms, with a 0.5ms rise and fall time. Stimuli were created in SigGen software (Tucker-Davis Technologies [TDT], Alachua, FL) and generated by a RZ6 multi-I/O processor (TDT). Stimuli were played from a free field speaker (MF1, TDT) located 10cm from the animal’s left pinna at 0° azimuth. Stimuli were calibrated with a 0.25 in. free-field microphone (PCB Piezotronics, Depew, NY, model 378C01) placed at the location of the animal’s pinna.

ABR traces were analyzed by two researchers independently, one completely blinded to the experimental conditions of the subjects. Inter-rater reliability was >98%. ABR threshold was defined as the average between the lowest sound level to evoke a response and the first level with no response (any wave). Peak-to-trough amplitudes were derived for ABR Waves I manually.

### Acoustic exposure protocol

Awake, unrestrained mice were exposed to a 100dB SPL octave-band (8–16kHz) noise for 1h. Mice were put in a custom-built soundproof chamber (53.5 × 54.5 × 57 cm) lined with 4cm-thick Sonex sound attenuating foam (Illbruck Inc.) within individual wire cells (10.5cm* 5cm* 5cm), where they moved freely. A Fostex dome tweeter speaker (FT28D), mounted atop a chicken wire bridge (0.5 × 0.5-inch squares) was suspended over the center of the mouse cage in the booth. The Fostex speaker was wired to an Amplifier (Crown D-75A), which was hooked up to a Dell Optiplex computer (Model 580). The sound file was 5 min in duration and was presented on a continuous loop for the duration of the noise exposure (1 hour). Calibrations were conducted using a Larson Davis sound level meter (System 824) to ensure the noise was 100dB (Z-weighted scale) in all four corners and in the center of the cage. Calibrations took place before every noise exposure.

### Immunohistochemistry and Quantification

After 14-day ABRs concluded, mice were euthanized and both cochleas fixed in 4% paraformaldehyde for 30 minutes at room temperature. The cochleas were decalcified in EDTA 250mM for 2-3 days at RT, following dissection, then underwent blocking for immunohistology. Primary antibodies used are Cell Signaling Technology HA-Tag (C29F4) Rabbit mAb #3724 (1:100), DSHB Mouse IgG1 anti-SV2 (1:1000), Biolegend Purified Mouse IgG2α,κ anti-PSD95 Cat # 810401 (Previously Covance catalog# MMS-5182) (1:200) and BD Transduction Laboratories Cat # 612044 Clone 16 Purified Mouse IgG1 anti-CtBP2 (1:200). Secondary AlexaFluor-conjugated antibodies from Invitrogen (1:1000) were used with DAPI (1:2000) to label the nuclei. Whole mount cochlear sections were mounted with ProLong Gold antifade reagent (Cat #P36930, Thermo Fisher Scientific). Images were acquired on an inverted Nikon A1 confocal microscope using 40× NA 1.3 and 63× NA 1.4 oil immersion objectives.ImageJ/Fiji (RRID: SCR_002285) was used for image analysis and counts. Synaptic puncta were manually counted by one observer.

### Statistical analysis

Mean± SEM are plotted. Data was analyzed with GraphPad Prism 10.2.3. Statistical analyses included ordinary one way ANOVA with Tukey’s multiple comparisons test (if normality present), Kruskal Wallis test (if non normal data) or Friedman’s test with Dunn’s multiple comparisons (if repeated measures) for clicks and two-way ANOVA with multiplicity adjusted P-values after Tukey’s multiple comparisons test or repeated measures two-way ANOVA with multiplicity adjusted P-values with either Dunnett’s or Tukey’s tests for pure tones. Two-way ANOVA was used for IHC synapse counts. Differences were considered statistically significant if P < 0.05.

## Acknowledgments

We thank F. Chakir for outstanding technical support, Drs. H. Hiel and A. Lauer for guidance on histology and ABRs, respectively.

## Funding

Support provided by R01 DC001508 and R01 DC016559 from the National Institutes for Deafness and Communication Disorders, NIH, by the David M. Rubenstein Professorship and Fund for Hearing Research at Johns Hopkins University School of Medicine, and by an Emerging Research Grant from the Hearing Health Foundation to MW.

## Author contributions

Data collection:

ES Analysis: ES, PAF, MW

Project direction: ES, PAF, MW

Funding Acquisition: PAF, MW

Writing - original draft: ES

Writing – Revisions & editing: ES, PAF, MBW

## Competing interests

Authors declare that they have no competing interests.

## Data and materials availability

All data including statistical validation, are included in the paper. Research materials are available via a Materials Transfer Agreement issued by the Johns Hopkins office of Tech Transfer.

## Notes

### Competing Interest Statement

The authors have declared no competing interest.

## References and Notes

1. Chadha, S., K. Kamenov, and A. Cieza, The world report on hearing, 2021. Bulletin of the World Health Organization, 2021/04/04. 99(4).

2. Collaborators, G.H.L., Hearing loss prevalence and years lived with disability, 1990–2019: findings from the Global Burden of Disease Study 2019. Lancet (London, England), 2021/03/03. 397(10278).

3. McDaid, D., A.-L. Park, and S. Chadha, Estimating the global costs of hearing loss. International Journal of Audiology, 2021-3-1. 60(3).

4. Global and regional needs, unmet needs and access to hearing aids - PubMed. International journal of audiology, 2020 Mar. 59(3).

5. Scholes, S., et al., Socioeconomic differences in hearing among middle-aged and older adults: cross-sectional analyses using the Health Survey for England. BMJ Open, 2018. 8(2).

6. Basner, M., et al., Auditory and non-auditory effects of noise on health. Lancet, 2014/04/04. 383(9925).

7. Fuchs, P.A. and A.M. Lauer, Efferent Inhibition of the Cochlea. Cold Spring Harbor Perspectives in Medicine, 2019/05. 9(5).

8. Warr, W.B. and J.J. Guinan, Jr., Efferent innervation of the organ of corti: two separate systems. Brain Res, 1979. 173(1): p. 152–5.

9. Roux, I., et al., Onset of Cholinergic Efferent Synaptic Function in Sensory Hair Cells of the Rat Cochlea. Journal of Neuroscience, 2011-10-19. 31(42).

10. Vattino, L.G., et al., Functional Postnatal Maturation of the Medial Olivocochlear Efferent–Outer Hair Cell Synapse. Journal of Neuroscience, 2020-06-17. 40(25).

11. Johnson, S.L., et al., Cholinergic efferent synaptic transmission regulates the maturation of auditory hair cell ribbon synapses. Open Biology, 2013/11. 3(11).

12. Synaptic specializations associated with the outer hair cells of the Japanese macaque - PubMed. Hearing research, 1997 Jun. 108(1-2).

13. Structure of the nerve endings on the external hair cells of the guinea pig cochlea as studied by serial sections - PubMed. Journal of ultrastructure research, 1961 Dec. 5(6).

14. Fuchs, P.A., M. Lehar, and H. Hiel, Ultrastructure of Cisternal Synapses on Outer Hair Cells of the Mouse Cochlea. The Journal of comparative neurology, 2014/02/02. 522(3).

15. Wersinger, E., et al., BK Channels Mediate Cholinergic Inhibition of High Frequency Cochlear Hair Cells. PLoS ONE, 2010. 5(11).

16. Elgoyhen, A.B. and E. Katz, The efferent medial olivocochlear-hair cell synapse. J Physiol Paris, 2012. 106(1-2): p. 47–56.

17. Boero, L.E., et al., Enhancement of the Medial Olivocochlear System Prevents Hidden Hearing Loss. J Neurosci, 2018. 38(34): p. 7440–7451.

18. Chumak, T., et al., Deterioration of the Medial Olivocochlear Efferent System Accelerates Age-Related Hearing Loss in Pax2-Isl1 Transgenic Mice. Molecular Neurobiology 2015 53:4, 2015-05-20. 53(4).

19. Galambos, R., SUPPRESSION OF AUDITORY NERVE ACTIVITY BY STIMULATION OF EFFERENT FIBERS TO COCHLEA. Journal of Neurophysiology, 1956 Sep 01. 19(5).

20. Kujawa, S.G. and M.C. Liberman, Conditioning-Related Protection From Acoustic Injury: Effects of Chronic Deefferentation and Sham Surgery. Journal of Neurophysiology, 1997 Dec 01. 78(6).

21. The olivocochlear efferent bundle and susceptibility of the inner ear to acoustic injury. Journal of Neurophysiology, 1991. 65(1).

22. Liberman, M.C., L.D. Liberman, and S.F. Maison, Efferent Feedback Slows Cochlear Aging. The Journal of Neuroscience, 2014/03/03. 34(13).

23. Maison, S.F. and M.C. Liberman, Predicting Vulnerability to Acoustic Injury with a Noninvasive Assay of Olivocochlear Reflex Strength. The Journal of Neuroscience, 2000/06/06. 20(12).

24. Maison, S.F., et al., Efferent Protection from Acoustic Injury Is Mediated via α9 Nicotinic Acetylcholine Receptors on Outer Hair Cells. The Journal of Neuroscience, 2002. 22(24): p. 10838–10846.

25. Maison, S.F., H. Usubuchi, and M.C. Liberman, Efferent Feedback Minimizes Cochlear Neuropathy from Moderate Noise Exposure. The Journal of Neuroscience, 2013/03/03. 33(13).

26. Reiter, E.R. and M.C. Liberman, Efferent-mediated protection from acoustic overexposure: relation to slow effects of olivocochlear stimulation. Journal of Neurophysiology, 1995 Feb 01. 73(2).

27. Wiederhold, M.L. and N.Y.S. Kiang, Effects of Electric Stimulation of the Crossed Olivocochlear Bundle on Single Auditory-Nerve Fibers in the Cat. The Journal of the Acoustical Society of America, 19701/10/01. 48(4B).

28. Zhang, Y., et al., Engineering olivocochlear inhibition to reduce acoustic trauma. Mol Ther Methods Clin Dev, 2023. 29: p. 17–31.

29. Taranda, J., et al., A point mutation in the hair cell nicotinic cholinergic receptor prolongs cochlear inhibition and enhances noise protection. PLoS Biol, 2009. 7(1): p. e18.

30. Isgrig, K., et al., AAV2.7m8 is a powerful viral vector for inner ear gene therapy. Nat Commun, 2019. 10(1): p. 427.

31. Vyas, P., et al., Characterization of HA-tagged α9 and α10 nAChRs in the mouse cochlea. Sci Rep, 2020. 10(1): p. 21814.

32. Talaei, S., et al., Dye Tracking Following Posterior Semicircular Canal or Round Window Membrane Injections Suggests a Role for the Cochlea Aqueduct in Modulating Distribution. Front Cell Neurosci, 2019. 13: p. 471.

33. Refining surgical techniques for efficient posterior semicircular canal gene delivery in the adult mammalian inner ear with minimal hearing loss - PubMed. Scientific reports, 09/22/2021. 11(1).

34. Stöver, T., M. Yagi, and Y. Raphael, Transduction of the contralateral ear after adenovirus-mediated cochlear gene transfer. Gene Ther, 2000. 7(5): p. 377–83.

35. Plontke, S.K. and A.N. Salt, Local drug delivery to the inner ear: Principles, practice, and future challenges. Hear Res, 2018. 368: p. 1–2.

36. Salt, A.N. and K. Hirose, Communication pathways to and from the inner ear and their contributions to drug delivery. Hear Res, 2018. 362: p. 25–37.

37. Hesse, L., et al., Non-Monotonic Relation Between Noise Exposure Severity and Neuronal Hyperactivity in the Auditory Midbrain. Frontiers in Neurology, 2016. 7.

38. Gittleman, S.N., C.G.L. Prell, and T.L. Hammill, Octave band noise exposure: Laboratory models and otoprotection efforts. The Journal of the Acoustical Society of America, 2019/11. 146(5).

39. Kujawa, S.G. and M.C. Liberman, Synaptopathy in the noise-exposed and aging cochlea: primary neural degeneration in acquired sensorineural hearing loss. Hearing research, 2015/12. 330(0 0).

40. Cheatham, M.A., et al., Cochlear function in Prestin knockout mice. The Journal of Physiology, 2004/11/11. 560(Pt 3).

41. Fernandez, K.A., et al., Noise-induced Cochlear Synaptopathy with and Without Sensory Cell Loss. Neuroscience, 2020/02/02. 427.

42. Kujawa, S.G. and M.C. Liberman, Adding Insult to Injury: Cochlear Nerve Degeneration after “Temporary” Noise-Induced Hearing Loss. The Journal of Neuroscience, 2009/11/11. 29(45).

43. Translational issues in cochlear synaptopathy. Hearing Research, 2017/06/01. 349.

44. Primary neural degeneration in the Guinea pig cochlea after reversible noise-induced threshold shift - PubMed. Journal of the Association for Research in Otolaryngology : JARO, 2011 Oct. 12(5).

45. Time course of cochlear injury discharge (excitotoxicity) determined by ABR monitoring of contralateral cochlear events - PubMed. Hearing research, 2014 Sep. 315.

46. Protection of cochlear synapses from noise-induced excitotoxic trauma by blockade of Ca2+-permeable AMPA receptors - PubMed. Proceedings of the National Academy of Sciences of the United States of America, 02/18/2020. 117(7).

47. Wagner, E.L. and J.-B. Shin, Mechanisms of hair cell damage and repair. Trends in neurosciences, 2019/06. 42(6).

48. Lv, J., et al., AAV1-hOTOF gene therapy for autosomal recessive deafness 9: a single-arm trial. The Lancet, 2024. 403(10441): p. 2317–2325.

49. Qi, J., et al., AAV-Mediated Gene Therapy Restores Hearing in Patients with DFNB9 Deafness. Advanced Science, 2024/03. 11(11).

50. Elgoyhen, A.B., The α9α10 nicotinic acetylcholine receptor: a compelling drug target for hearing loss? Expert Opinion on Therapeutic Targets, 2022-03-04. 26(3).

51. Orthosteric and allosteric potentiation of heteromeric neuronal nicotinic acetylcholine receptors. British Journal of Pharmacology, 2018. 175(11).

52. Gu, S., et al., Hair cell α9α10 nicotinic acetylcholine receptor functional expression regulated by ligand binding and deafness gene products. Proceedings of the National Academy of Sciences of the United States of America, 2020/09/09. 117(39).

53. Morley, B.J., P. Whiteaker, and A.B. Elgoyhen, Commentary: Nicotinic Acetylcholine Receptor α9 and α10 Subunits Are Expressed in the Brain of Mice. Frontiers in Cellular Neuroscience, 2018. 12.

54. Christensen, S.B., et al., RgIA4 Potently Blocks Mouse α9α10 nAChRs and Provides Long Lasting Protection against Oxaliplatin-Induced Cold Allodynia. Frontiers in Cellular Neuroscience, 2017. 11.

55. Hone, A.J. and J.M. McIntosh, Nicotinic Acetylcholine Receptors in Neuropathic and Inflammatory Pain. FEBS letters, 2018/04. 592(7).

56. Hone, A.J., D. Servent, and J.M. McIntosh, α9-containing nicotinic acetylcholine receptors and the modulation of pain. British Journal of Pharmacology, 2018/06. 175(11).

57. Nowak, N., et al., Prior Acoustic Trauma Alters Type II Afferent Activity in the Mouse Cochlea. eNeuro, >2021-11-01. 8(6).

58. Wood, M.B., N. Nowak, and P.A. Fuchs, Frontiers | Damage-evoked signals in cochlear neurons and supporting cells. Frontiers in Neurology, 2024/02/14. 15.

59. Unmyelinated type II afferent neurons report cochlear damage - PubMed. Proceedings of the National Academy of Sciences of the United States of America, 11/24/2015. 112(47).

60. Outer Hair Cell Glutamate Signaling through Type II Spiral Ganglion Afferents Activates Neurons in the Cochlear Nucleus in Response to Nondamaging Sounds - PubMed. The Journal of neuroscience : the official journal of the Society for Neuroscience, 03/31/2021. 41(13).

61. Dalkara, D., et al., In Vivo–Directed Evolution of a New Adeno-Associated Virus for Therapeutic Outer Retinal Gene Delivery from the Vitreous. Science Translational Medicine, 2013-06-12. 5(189).

